# Distinct oscillatory frequencies underlie excitability of human occipital and parietal cortex

**DOI:** 10.1101/082693

**Authors:** Jason Samaha, Olivia Gosseries, Bradley R. Postle

## Abstract

Magnetic stimulation (TMS) of human occipital and posterior parietal cortex can give rise to visual sensations called phosphenes, but neural correlates of phosphene perception preceding and succeeding stimulation of both areas are unknown. Using near-threshold TMS with concurrent electroencephalography (EEG) recordings, we uncover oscillatory brain dynamics that covary, on single trials, with the perception of phosphenes following occipital and parietal TMS. Prestimulus power and phase predominantly in the alpha-band (8-13 Hz) predicted occipital TMS phosphenes, whereas higher frequency beta-band (13-20 Hz) power (but not phase) predicted parietal TMS phosphenes. TMS *evoked* responses related to phosphene perception were similar across stimulation sites and were characterized by an early (200 ms) posterior negativity and a later (>300 ms) parietal positivity in the time domain and an increase in low-frequency (~5-7 Hz) power followed by a broadband decrease in alpha/beta power in the time-frequency domain. These correlates of phosphene perception closely resemble known electrophysiological correlates of conscious perception using near-threshold visual stimuli and speak to the possible early onset of visual consciousness. The differential pattern of prestimulus predictors of phosphene perception suggest that distinct frequencies reflect cortical excitability within different cortical regions, and that the alpha-band rhythm, long thought of as a general index of cortical inhibition, may not reflect excitability of posterior parietal cortex.

**Significance statement:** Alpha-band oscillations are thought to reflect cortical excitability and are therefor suggested to play an important role in gating information transmission across cortex. We directly probe cortical excitability in human occipital and parietal cortex and observed that whereas alpha-band dynamics indeed reflect excitability of occipital areas, beta-band activity was most predictive of parietal cortex excitability. Differences in the state of cortical excitability predicted perceptual outcomes, which were manifest in both early and late patterns of evoked activity, shedding light on the neural correlates of consciousness. Our findings prompt revision of the notion that alpha activity reflects inhibition across all of cortex and suggests instead that excitability in different regions is reflected in distinct frequency bands.

## Introduction

Transcranial magnetic stimulation (TMS) can elicit muscle contractions when applied over motor cortex (1) and can induce visual sensations (phosphenes) when applied over regions of occipital cortex (2). These phenomena can be used to study the state of cortical excitability of the tissue undergoing stimulation by measuring changes in the magnitude or likelihood of these responses as stimulation parameters are kept constant (3). When TMS is combined with concurrent electroencephalographic (EEG) recordings, variability in cortical excitability can be linked to variability in ongoing neural activity. Using this approach, prior work has found that the power of oscillatory neural activity in the alpha band (8-13 Hz), prior to the onset of TMS, is negatively correlated with both phosphene perception (4) and the likelihood and magnitude of motor-evoked potentials (5, 6). These findings are taken as evidence that cortical excitability is regulated by neural activity in the alpha range (7). (Although prestimulus beta-band oscillations between 13 and 30 Hz also predict the amplitude of motor evoked potentials (8, 9)). Recent studies have also found that the phase of alpha-band oscillations is predictive of phosphene perception (10) as well as detection and discrimination of visual stimuli (11–13), leading to the proposal that alpha may impose inhibition in a phasic manner (14).

Because using this TMS-EEG approach to probe cortical excitability relies on stimulating brain areas that give rise to observable or reportable responses (e.g., phosphenes), previous work has been limited to investigating excitability of visual and motor cortex. Recently, however, several groups have reported that TMS applied to posterior parietal cortex (PPC) can also give rise to phosphenes (15–18), the phenomenology of which is comparable to those elicited from occipital stimulation (19). We therefore sought to characterize cortical excitability of PPC as well as occipital cortex by applying TMS at phosphene-threshold levels to both areas while recording EEG. Our approach further allowed us to investigate the neural correlates of phosphene perception in the TMS-evoked responses. Although prior work has investigated event-related potential (ERP) correlates of visual awareness using near-threshold visual stimuli (reviewed in (20, 21) as well as phosphene induction (17, 22), ERPs capture a limited amount of information about neural activity (only the phase-locked component) and no study to date has reported on oscillatory correlates of phosphene perception following occipital and parietal TMS.

Prior work investigating ERP correlates of conscious visual perception has identified two ERP components that vary with awareness. The so-called “visual-awareness negativity” (VAN) is an early component (~200 ms) that shows larger negative-going amplitudes at central- and lateral-occipital sites to consciously perceived stimuli across a wide array of awareness manipulations (23–28). The late positive potential (LP) occurs after 300 ms at central-parietal sites and is larger in amplitude to consciously perceived stimuli (29, 30). As these two components differ with respect to both the temporal onset and the cortical localization of conscious perception, debate is ongoing as to which pattern of activity reflects the neural correlate of perceptual awareness (31, 32).

Regarding the prestimulus predictors of occipital TMS phosphene perception, we sought to replicate prior findings of an inverse relationship between alpha power over posterior electrodes and subsequent phosphene visibility (4) as well as a dependence of phosphene perception on prestimulus alpha phase (10). Regarding parietal TMS phosphenes, it may be the case that alpha dynamics reflect cortical excitability outside of primary visual and motor cortices, in which case alpha power and/or phase may also predict parietal TMS phosphenes. Alternatively, given the prominent role of beta-band oscillations in parietal cortex for perceptual decision making (33, 34), activity in higher frequencies may be predictive of parietal TMS phosphenes. Data in favor of this view come from a recent experiment where TMS was applied to occipital, posterior parietal, and frontal cortex while EEG was recorded. The researchers observed that the dominant frequency of the TMS-evoked response increased from ~10 Hz following occipital stimulation, to ~ 18-20 Hz following parietal TMS, to ~ 31 Hz following frontal TMS (35). This finding was later replicated (36), and was taken to suggest that different cortical regions are intrinsically tuned to generate oscillations at different frequencies, with alpha dominating in occipital cortex and beta dominating in PPC. In line with this suggestion, prestimulus dynamics around 20 Hz were found to predict the global mean field amplitude of the response to PPC TMS (37).

Here, we stimulate occipital and posterior parietal cortex while recording EEG. We measured phosphene visibility reports on a continuous scale, which allowed us to capture the gradedness of perceptual experience, as suggested by prior work (38, 39), and which served as a measurement for single-trial regression analyses relating phosphene visibility to ongoing fluctuations in EEG voltage, power, and phase. We observed negative correlations between phosphene visibility and prestimulus low-frequency theta/alpha rhythms during occipital TMS and found negative correlations with prestimulus beta-band oscillations during parietal TMS. Phosphene perception following occipital TMS was also dependent on prestimulus alpha phase, whereas no phase dependency was found for parietal TMS phosphenes. Poststimulus correlates of both phosphene types were highly similar in timing and frequency characteristics, although early voltage effects had noticeably different scalp topographies. These findings provide evidence regarding the precise onset and oscillatory correlates of occipital and parietal TMS-evoked phosphene perception and suggest that alpha-band oscillations reflect cortical excitability of occipital, but not posterior parietal, cortex which may instead be related to beta oscillations.

## Materials and Methods

### Subjects

17 subjects were recruited for the experiment for monetary compensation. Following phosphene screening, which entailed single-pulse stimulation of right occipital and right parietal cortex up to 90% of maximum stimulator output, 10 subjects (8 male; 24-33 years old) who perceived phosphenes reliably at both stimulation sites were retained for EEG recording. This proportion is consistent with reports of 60-80% of subjects who perceive occipital TMS phosphenes without extensive training and resulted in a sample size that is comparable to previous TMS-EEG experiments examining phosphenes (4, 10, 17). All subjects were recruited from the University of Wisconsin-Madison community. The UW-Madison Health Sciences Institutional Review Board approved the study protocol. All subjects gave informed consent and were screened for the presence of neurological and psychiatric conditions and other risk factors related to the application of TMS.

### Stimulation and phosphene thresholding

Following others (15–19), stimulation of right occipital cortex was delivered over electrode O2 and stimulation of right PPC was delivered over electrode P4. The coil handle was oriented medial to lateral, away from the inion, during occipital stimulation and ventromedial to dorsolateral, pointing away from the inion for parietal stimulation (see arrows in Figure 1B). The final coordinates of stimulation were then determined functionally by slightly adjusting the coil until phosphenes were reliably elicited. Phosphene perception was considered reliable if participants reported them in the visual field contralateral to stimulation and if they ceased with decreased stimulation intensity. To ensure consistent stimulation throughout the experiment, the final stimulation coordinates were saved and visualized using a Navigated Brain Stimulation (NBS) system (Nextstim, Helsinki, Finland) that uses infrared-based frameless stereotaxy to map the position of the coil and the subject’s head within the reference space of the individual’s high-resolution T1-weighted anatomical MRI (acquired with a GE MR750 3-T scanner for each subject prior to the experiment). During the EEG recording the coil was held in place by the experimenter who monitored the NBS system to maintain consistent stimulation coordinates. Stimulator intensity was determined for each subject and stimulation site prior to the experiment as the intensity that lead to a 50% rate of phosphene reports following 10 single-pulse stimulations. As was also noted in prior work (16, 17, 19), parietal TMS phosphenes thresholds required higher stimulator intensities (*M* = 83.2% of maximum stimulator output SD = 9.2) than occipital (*M* = 71.9% SD = 8.1). A thin layer of foam was placed between the coil and the EEG cap to help minimize TMS artifacts and auditory artifacts due to bone conduction (40). This resulted in a higher intensity stimulator output, as compared to prior work, in order to achieve phosphene thresholds. TMS was delivered with a Magstim Super Rapid^2^ magnetic stimulator fit with a focal bipulse, figure of eight 70-mm stimulating coil (Magstim, Whitland, UK).

**Figure 1.**
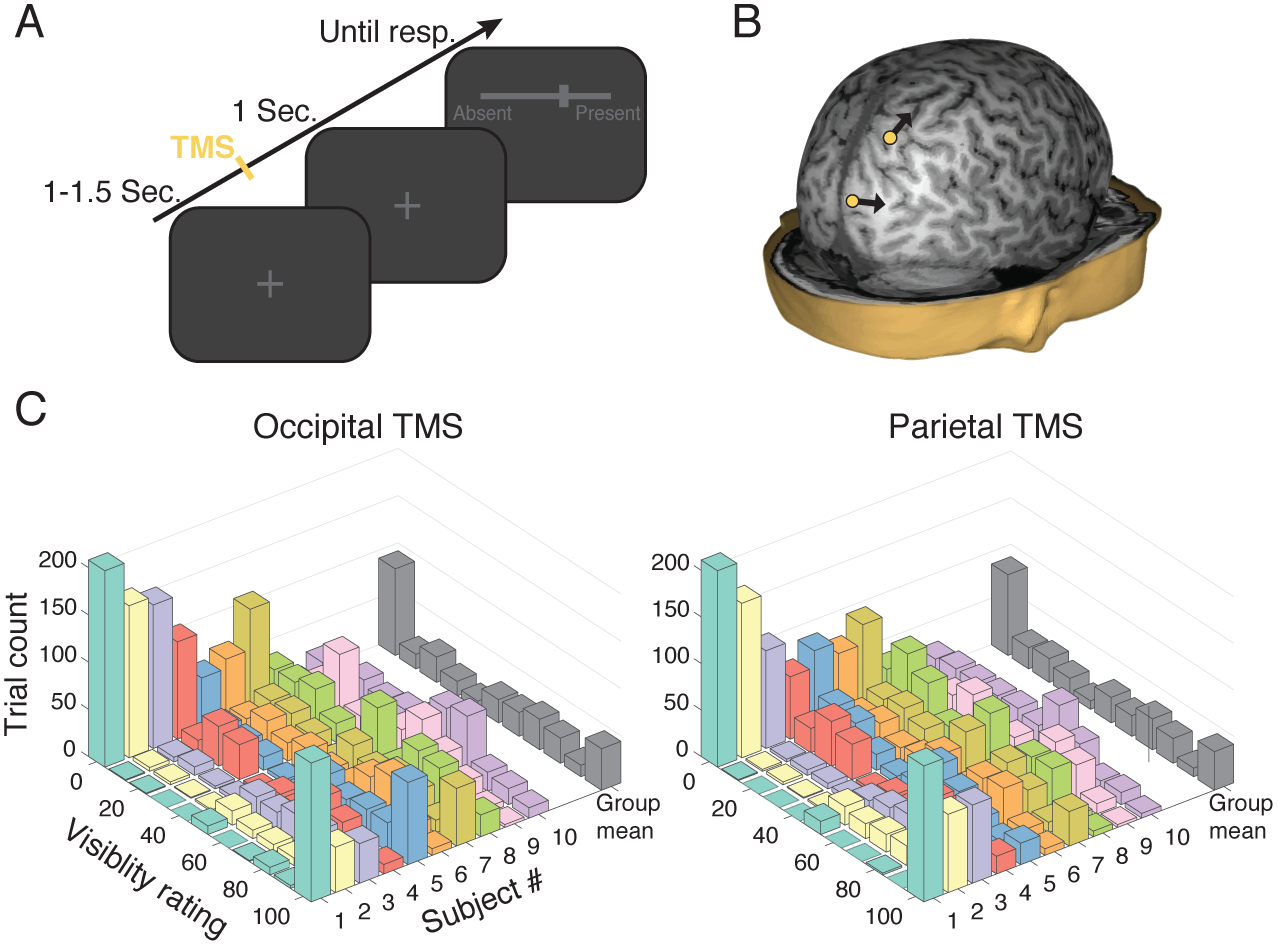
Task design, stimulation sites, and visibility response distributions. (A). Following a variable inter-trial interval of 1-1.5 seconds, a single pulse of TMS was delivered at individual phosphene thresholds. One second after TMS, subjects reported the visibility of their phosphene perception by sliding a cursor across a one-dimensional scale between “clearly absent” and “clearly present”. (B) Stimulation was applied to functionally determined targets within occipital and parietal cortex in counterbalanced blocks. The yellow dots indicate targets from a representative subject and the arrows indicate the orientation of the handle of the TMS coil. (C) Individual subject and group level visibility distributions following occipital and parietal TMS.

### Experimental session

Each subject completed 8 blocks of 100 trials each. Single-pulse TMS was delivered to occipital cortex on half of the blocks and to parietal cortex on the remaining half. Occipital and parietal stimulation blocks were interleaved and the order was counterbalanced across participants. The task is depicted in Figure 1A. Each trial began with a random inter-trial interval between 1000 and 1500 ms during which a dark grey fixation cross was presented atop a black background. Following each pulse of TMS, subjects’ maintained central fixation for 1 second at which point a dark grey scale appeared. The left and right endpoints of the semi-continuous scale (0-100) were respectively labeled “clearly absent” and “clearly present”. Subjects were instructed to rate the visibility of their phosphene perception by sliding a computer mouse with their right hand to the appropriate position on the scale and clicking the mouse. The current scale position was marked by a perpendicular bar and was reset to the center of the scale on each trial. As a check of basic task compliance, TMS was not delivered on 5% of all trials (randomly determined), which we expected would result in reports on the lower end of the visibility scale.

### EEG acquisition and preprocessing

EEG was recorded from 60 Ag/AgCl electrodes connected to a TMS-compatible amplifier (Nexstim, Helsinki, Finland). This amplifier avoids saturation by the TMS pulse with a sample-and-hold circuit that holds amplifier output constant from 100 µs before to 2 ms after stimulus. Impedance at each electrode was kept below 5 kΩ. A single electrode placed on the forehead was used as the reference, and eye movements were recorded with two additional electrodes placed near the eyes. Data were acquired at a rate of 1,450 Hz with 16-bit resolution. To reduce contamination of the EEG by auditory responses from TMS, masking white noise was played through inserted earplugs throughout the experimental session, as in previous experiments in our lab (37). EEG was processed offline with custom MATLAB scripts (version R2014b) and with the EEGLAB toolbox version 13.5 (41). Recordings were visually inspected and noisy channels (3.9 on average) were spherically interpolated. Data were then downsampled to 500 Hz and re-referenced to the average of all electrodes. Because for some subjects residual high-amplitude artifacts persisted for 20-30 ms following TMS, the data from -10 to 40 ms surrounding each TMS pulse was removed and interpolated via robust splines (42). A one-pass zero-phase Hamming windowed-sinc FIR filter between 0.5 and 50 Hz was applied to the data (EEGLAB function *pop_eegfiltnew.m*) and epochs spanning -1500 to 1500 relative to TMS onset were extracted. A prestimulus baseline of -200 to -10 was then subtracted from each trial. Individual trials were then visually inspected and those containing muscle artifacts or ocular artifacts occurring contemporaneously with TMS onset were removed, resulting in an average of 322 occipital stimulation and 323 parietal stimulation trials remaining per subject. Independent components analysis using the INFOMAX algorithm (EEGLAB function *binica.m*) was used to remove remaining ocular artifacts not coinciding with TMS as well as artifactual components clearly related to TMS. An average of 2.7 ocular artifacts and 6.8 TMS-related artifacts were removed per subject. Raw data and commented code used for all preprocessing and analysis are freely available for download at the Open Science Framework (osf.io/6qu3b).

### Time-domain analysis

To relate continuously varying visibility ratings to continuously varying voltage across time, we performed non-parametric robust regression on single-trial data. For each time-point, electrode, and subject, regression coefficients that describe the monotonic relationship between voltage and visibility were estimated according to the linear model:

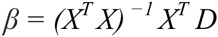

where *X* is the design matrix containing one column for the intercept (all ones) and one column for visibility ratings across trials and *^T^* and ^-1^ indicate the matrix transpose and inverse, and *D* is the vector of voltage data from all trials at a given time-point. The resulting beta coefficient representing the slope of the voltage-visibility relationship was then converted into a *z*-statistic relative to a subject-specific null hypothesis distribution obtained by repeatedly shuffling the mapping between visibility ratings and voltage (see *Statistics*). Prior to regression, the voltage data were smoothed with a 20 ms sliding-average window and both visibility and voltage were rank-scored to mitigate the influence of outlying data while testing for a monotonic relationship (this is equivalent to computing a Spearman’s correlation coefficient). These normalized beta coefficients were then averaged over a cluster of posterior electrodes (visualized in the Figure 2B inset) to improve signal-to-noise. To validate this procedure against a more traditional time-domain analysis approach, trial-averaged ERPs were also computed by sorting trials into “high” and “low” visibility bins if they were greater than (less than) the 55^th^ (45^th^) percentile of the visibility scale (in order to exclude middle “unsure” trials). Following others (24, 25, 28), the LP/P3 potential was examined at central-parietal electrode Pz and the VAN was examined at occipital electrode Oz. Time-windows for each component were determined from data orthogonal to later statistical contrasts by inspecting the condition-averaged ERP. VAN was averaged over a window that spanned 180-220 ms and the LP window spanned 300-800 ms.

**Figure 2.**
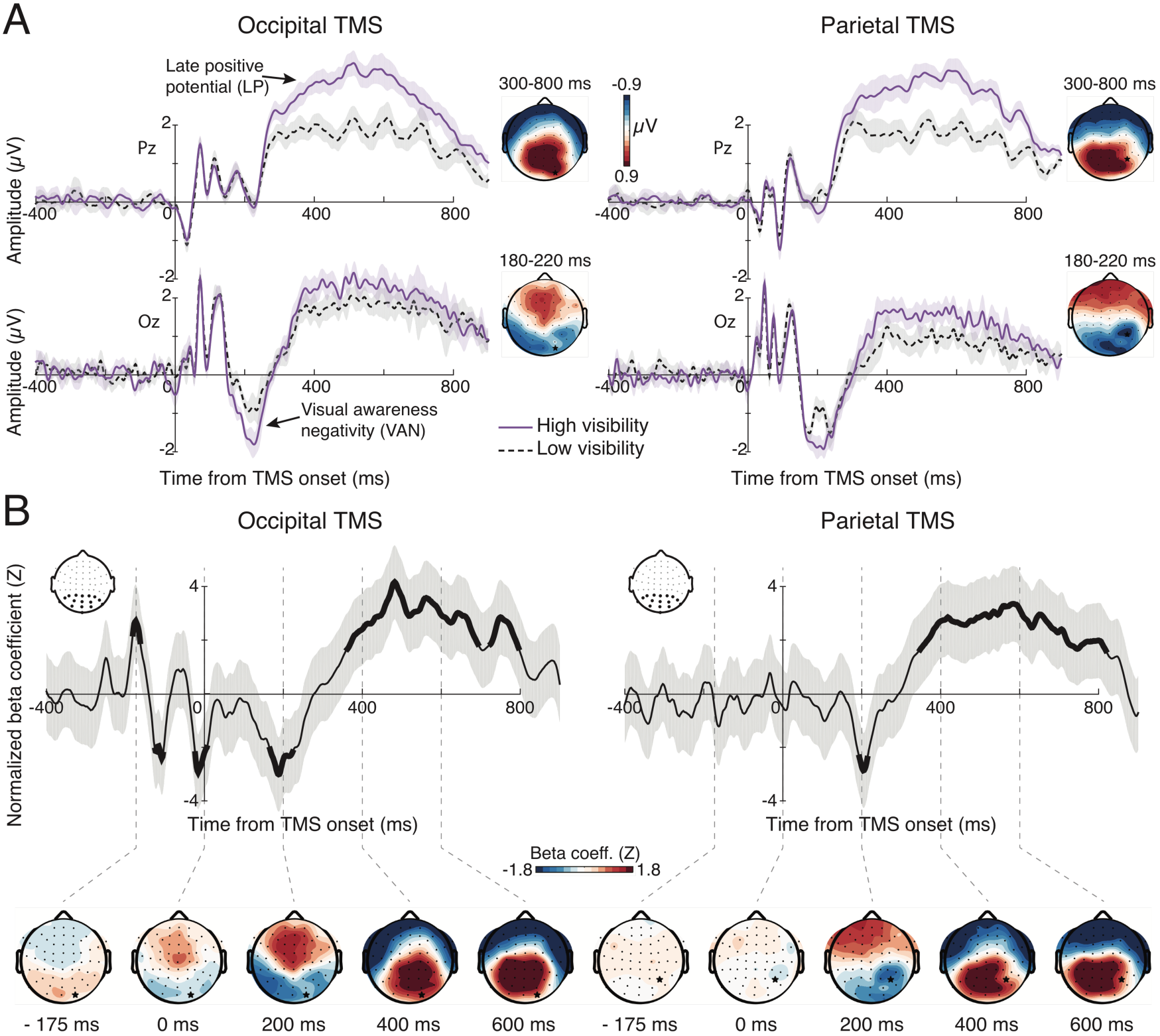
Voltage-domain correlates of phosphene perception time-locked to TMS onset. (A) ERPs contrasting high and low visibility for each stimulation condition revealed an early modulation of the VAN component (bottom) followed by modulation of the late positive potential (top). Scalp maps denote the difference topography of each component, revealing VAN modulation at lateral-occipital scalp sites following occipital TMS, and at central occipital-parietal sites following parietal TMS. The LPP modulation was parietally maximal at both stimulation sites (denoted with a black star). Shaded bands denote ±1 within-subject SEM (B) Single-trial regression results demonstrating the negative relation between voltage at 200 ms and visibility for both stimulation sites, and a positive relation between voltage after 300 ms and visibility. Note the similarity of the beta coefficient topographies to those obtained from the ERP analysis. Also of note is a strong prestimulus oscillation in beta coefficients prior to occipital TMS with a period of about 100 ms. This suggests an influence of prestimulus alpha-band phase on phosphene perception during occipital TMS. Thick line segments denote significant cluster-corrected regression coefficients and shaded bands represent 95% confidence intervals.

### Time-frequency analysis

The same single-trial regression approach was used to relate time-frequency power to visibility. Time-frequency decomposition was performed by convolving data from each trial with a family of complex Morlet wavelets spanning 2-50 Hz in 1.23-Hz steps with wavelet cycles increasing linearly between three and eight cycles as a function of frequency. Power was obtained by squaring the absolute value of the resulting complex time series and was converted to percent signal change relative to a prestimulus baseline of -600 to -100 ms in order to adjust for power-law scaling. Following regression, normalized beta coefficients were then averaged over the same cluster of posterior electrodes that was used in the time-domain regression analysis (above). Due to the temporal smearing inherent in time-frequency decomposition, caution must be used when analyzing prestimulus effects—particular with respect to phase (43)—which can result from a confound due to contamination of prestimulus data by poststimulus differences. Therefore, we focused further on prestimulus power and phase by performing an FFT on data segments cut from -1000 to -50 ms prior to TMS onset. Prior to the FFT, these segments were linearly detrended, multiplied by a Hamming-window, and zero-padded (frequency resolution: 0.1 Hz). Power was extracted by squaring the absolute value of the Fourier coefficients and phase was obtained by taking the angle (MATLAB function *angle.m*). Single-trial prestimulus power was related to visibility as before and was also binned into high and low visibility conditions to, again, illustrate differences in a more traditional manner. For this analysis, power was log_10_ transformed and averaged over the alpha-band (8-13 Hz) and low beta-band (13-20 Hz).

Because phase is a circular variable, it cannot be related to a linear variable by means of ordinary linear regression. We therefore computed a recently introduce measure of circular-linear association called *weighted inter-trial phase clustering* (wITPC; (44–46) to relate prestimulus phase to visibility ratings. wITPC is computed as the resultant vector length, or inter-trial phase clustering, (also called the phase locking factor or inter-trial coherence) of phase angles across trials once the length of each vector has been weighted by the linear variable of interest (an example of this computation with data from one subject is shown in Figure 4B). wITPC was computed for each subject, TMS condition, electrode, and frequency by multiplying the unit-length complex-valued phase angle on each trial by the corresponding trial’s visibility rating and then averaging those complex numbers across trials and taking the absolute value to obtain the resultant vector length. Because wITPC will be non-normally distributed and the magnitude will be strongly determined by the scale of the linear (weighting) variable, it is necessary to normalize this quantity with respect to a null distribution obtained by shuffling trial labels. Positive normalized wITPC values indicate that phase modulates visibility, or, in other words, that certain visibility ratings are more likely than chance to occur at certain phase angles of the measured oscillation.

**Figure 3.**
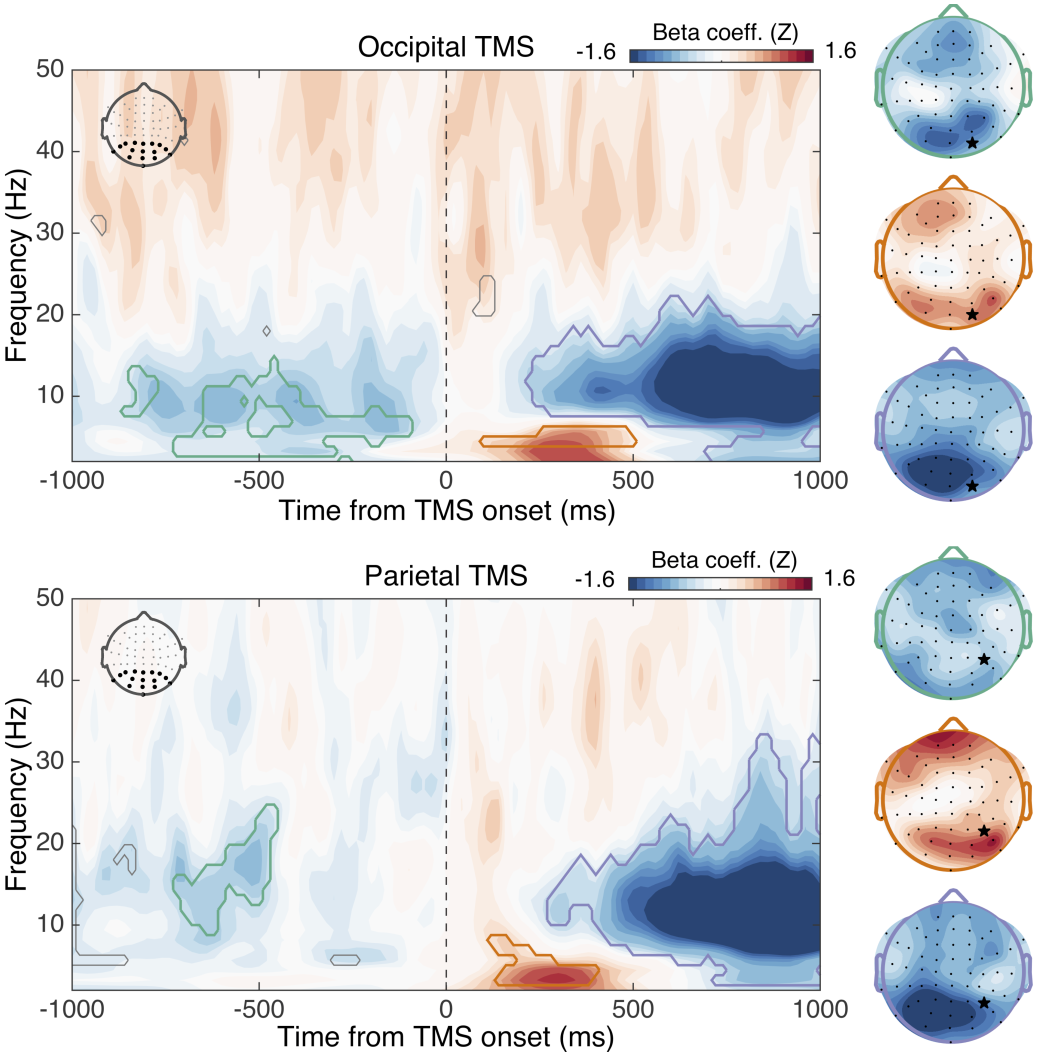
Time-frequency power correlates of phosphene perception. Maps show regression coefficients relating power at each time-frequency point to visibility ratings during occipital (upper panel) and parietal (lower panel) TMS. Contour lines encompass significant cluster-corrected effects and are color-coded according to their corresponding topography, displayed on the right side of the figure. Notably, poststimulus correlates of phosphene perception were highly similar across stimulation conditions, whereas prestimulus low frequency power (3-13 Hz) predicted phosphenes following occipital TMS and prestimulus beta-band power (10-22 Hz) predicted parietal TMS phosphenes. Note that the prestimulus scalp topographies (green outline) are displayed on a scale of ± 0.8. Star denotes stimulation site.

**Figure 4.**
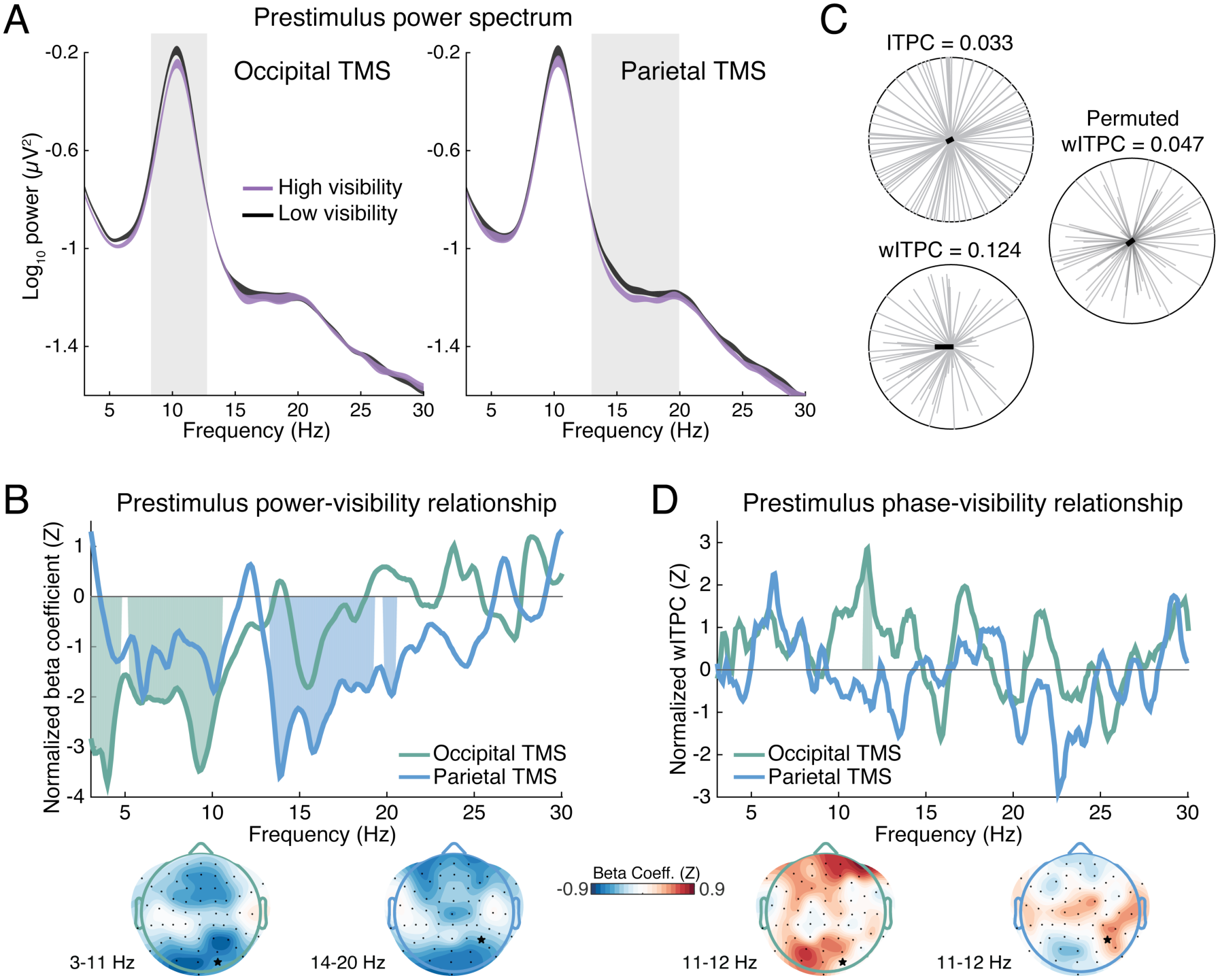
Relationship of prestimulus power and phase with visiblity. (A) Prestimulus power spectra showing modulation of alpha-band power prior to occipital tms and beta-band powerprior to parietal tms as a function of visiblity. Gray windows highlight frequency-bands of interest for statistical analysis. Bands are ±1 within-subject SEM. (B) This same pattern was borne out of a single-trial regression analysis on prestimulus power, demonstrating negative correlations between low-frequency power and occpital TMS phosphene visiblity (green line) and between beta-band power and parietal TMS phosphene visiblity (blue line). The topographies of both effects are maximal over poserior and fronal sensors. (C) Example computation of wITPC to relate single-trial phase to visibility. Upper left; single-trial prestimulus phase vectors are shown as gray lines and are not clustered across trials due to the temporal randomization of the inter-trial interval, leading to a low resultant vector length (i.e., low ITPC). Bottome left; the length of each trials phase vector is then weighted by that trial’s visiblity rating (here normalized between 0 and 1) and a weighted ITPC is computed, reflecting the degree of phase-visiblity correlation. Right; this quantitiy gets normalized with respect to a nule distribution attained by shuffling visiblity ratings across trials. (D). Computing normalized wITPC across frqeuencies revealed that prestimulus phase in the alpha-band (~11.6 Hz) was predictive of phosphene visiblity during occipital TMS. No phase-visiblity relantionship was found for parietal TMS.

### Statistics

Level one (subject-level) statistics were performed for all time and time-frequency domain single-trial analysis described above by randomly permuting the mapping between visibility ratings and neural data 1000 times, each time recomputing the relevant statistic (beta coefficient or wITPC). The statistic associated with the true data mapping was then converted to a z-statistic relative to the mean and standard deviation of the permuted data. This resulted in a z value for each analysis, subject, TMS condition, channel, and time- or time-frequency point. This approach incorporates knowledge about variability in the subject-level effects into the subsequent group-level analyses. Level two (group-level) statistics and significance values were also computed by means of non-parametric permutation tests in combination with threshold free cluster enhancement (TFCE) to address multiple comparisons across time and frequency points. To estimate group-level null hypothesis distributions, on each of 5000 permutations, z-scores from a random subset of subjects were multiplied by -1 and a t-test against zero was computed (this is equivalent to randomly swapping the order of the condition subtraction, e.g., A-B vs. B-A (47)). The t-statistic resulting from the true data mapping was then subject to TFCE as implemented in the LIMO EEG package (48), which uses the algorithm developed in (49). Each permutated t-statistic was also submitted to TFCE, forming a distribution of cluster extents expected under the null hypothesis. Only cluster extents in the real data exceeding an α rate of 0.05% were considered statistically significant. This procedure has been shown to control well the family-wise error rate across multiple comparisons while taking into account autocorrelation present in electrophysiological data (50). Paired t-tests were used to compare high versus low visibility ERP amplitudes and band-averaged FFT power. All tests were two-tailed.

## Results

### Visibility ratings

The average visibility rating (± SD) was 42.5 ± 8.9 (out of 100) following occipital stimulation and was 41.8 ± 4.8 following parietal stimulation, confirming that our thresholding procedure was effective. As shown in Figure 1C, subjects made use of the full range of the scale although with varying degrees of gradedness; some subjects used the endpoint values more frequently than others. When sorted into 11 bins (as in Figure 1C), the shape of the group-averaged histogram had 4 local maxima that were the same for occipital and parietal TMS. These peaks were at ratings of 4.5, 22.7, 68.18, and 95.5, consistent with previous work that has identified 4 “categories” of perceptual experience underlying the use of continuous visibility scales (51, 52). As expected, ratings on no-TMS trials were near zero (1.9 ± 3). These results suggest that additional information can be gained by allowing subjective visibility responses to take a non-binary form.

### Time-domain correlates of phosphene perception

ERP’s contrasting high and low visibility are presented in Figure 2A. High awareness trials were associated with larger TMS-evoked VAN amplitudes following both occipital (t(1,9) = -4.39, p = 0.002) and parietal (t(1,9) = -3.79, p = 0.004) TMS. This effect was similar across stimulation conditions, but had a more temporal-occipital scalp distribution in the occipital TMS condition and more central occipital-parietal distribution in the parietal TMS condition. High awareness was also associated with an enhanced LP potential over posterior electrodes for occipital (t(1,9) = 4.68, p = 0.001) and parietal (t(1,9) = 5.49, p < 0.001) TMS. These results were also borne out of the single-trial regression analysis (Figure 2B). In addition to significant negative correlations in the VAN time range (180-220 ms) and positive correlations in the LP time range (300-800), the single trial analyses revealed a robust prestimulus correlation during the occipital TMS condition. This prestimulus correlation had a reversing polarity over time, with a period of ~ 100 ms, suggesting that prestimulus alpha-band phase influenced phosphene visibility. The scalp distributions of these significant regression parameters highly resemble those attained from the ERP analysis, with the early (200 ms) visibility effect being maximal over occipital-temporal electrodes following occipital TMS and maximal over central occipital-parietal electrodes following parietal TMS and with the late effect being parietal-maximal in both stimulation conditions.

### Time-frequency power correlates of phosphene perception

The relationship between single-trial visibility and oscillatory power across time and frequency space is shown in Figure 3. The analysis of occipital TMS data revealed a negative relationship between prestimulus low-frequency power (~5-13 Hz) and visibility that had a posterior scalp distribution. TMS-evoked low-frequency power (~3-6 Hz) between ~140 and 400 ms over lateral occipital electrodes was positively correlated with visibility, whereas later (~250-900 ms) posterior alpha/beta power was robustly negatively correlated with visibility. Regarding parietal TMS, no clear effect of prestimulus low-frequency power was observed; rather, prestimulus high-alpha/low-beta power (10-24 Hz) was negatively related to phosphene visibility. The topography of this effect was more widespread, with both a posterior and a central-frontal distribution. The relationship between TMS-evoked power and phosphene visibility following parietal TMS closely resembled that of occipital stimulation: early low frequency power was positively predictive of visibility, followed by a broadband alpha/beta component that was negatively related to visibility.

### Prestimulus oscillatory power predicts phosphene perception

Figure 4A shows a complementary analysis of power focused just on the prestimulus interval, to avoid any contamination from poststimulus, TMS-induced responses (43). Power spectra of prestimulus data from high and low visibility trials show a clear alpha-band peak whose power was higher on low visibility occipital TMS trials (t(1,9) = -3.82, p = 0.004). In contrast, during parietal TMS, pre-stimulus beta power was significantly lower on high-visibility trials (t(1,9) = -2.92, p = 0.017). The results of the single-trial regression analysis (Figure 4B) revealed a clear distinction between the prestimulus predictors of phosphenes following occipital and parietal TMS, with broad-band low frequency power (3-13 Hz) negatively correlating with occipital TMS phosphene visibility and higher frequency beta-band (12-22 Hz) power negatively relating to parietal TMS phosphene visibility. The topographies of these regression coefficients closely resembled those obtained from the time-frequency described analysis above, both having maximal effects over posterior as well as frontal sensors.

### Prestimulus alpha phase predicts phosphenes following occipital TMS

Figure 4C depicts an example of wITPC computation (46) using a single subject’s prestimulus alpha phase during occipital TMS to predict visibility (see *Time-frequency analysis*, for more details). The results of this procedure applied across frequencies revealed a significant phase-visibility relationship in the alpha band (peak at 11.6 Hz) only during occipital TMS. This effect was maximal over both occipital and frontal electrodes. No significant phase-visibility effects survived correction for multiple comparisons during parietal TMS. The effect of prestimulus alpha phase on phosphene visibility was further demonstrated by binning trials into 7 bins according to prestimulus phase at 11.6 Hz (normalized to % change relative to the mean across all bins). As shown in Figure 5, this revealed a clear modulation of visibility according to prestimulus phase during occipital (F(6,54) = 3.23, p = 0.008), but not parietal TMS (F(6,54) = 1.88, p = 0.102), as determined from a oneway ANOVA predicting visibility from phase bin.

**Figure 5.**
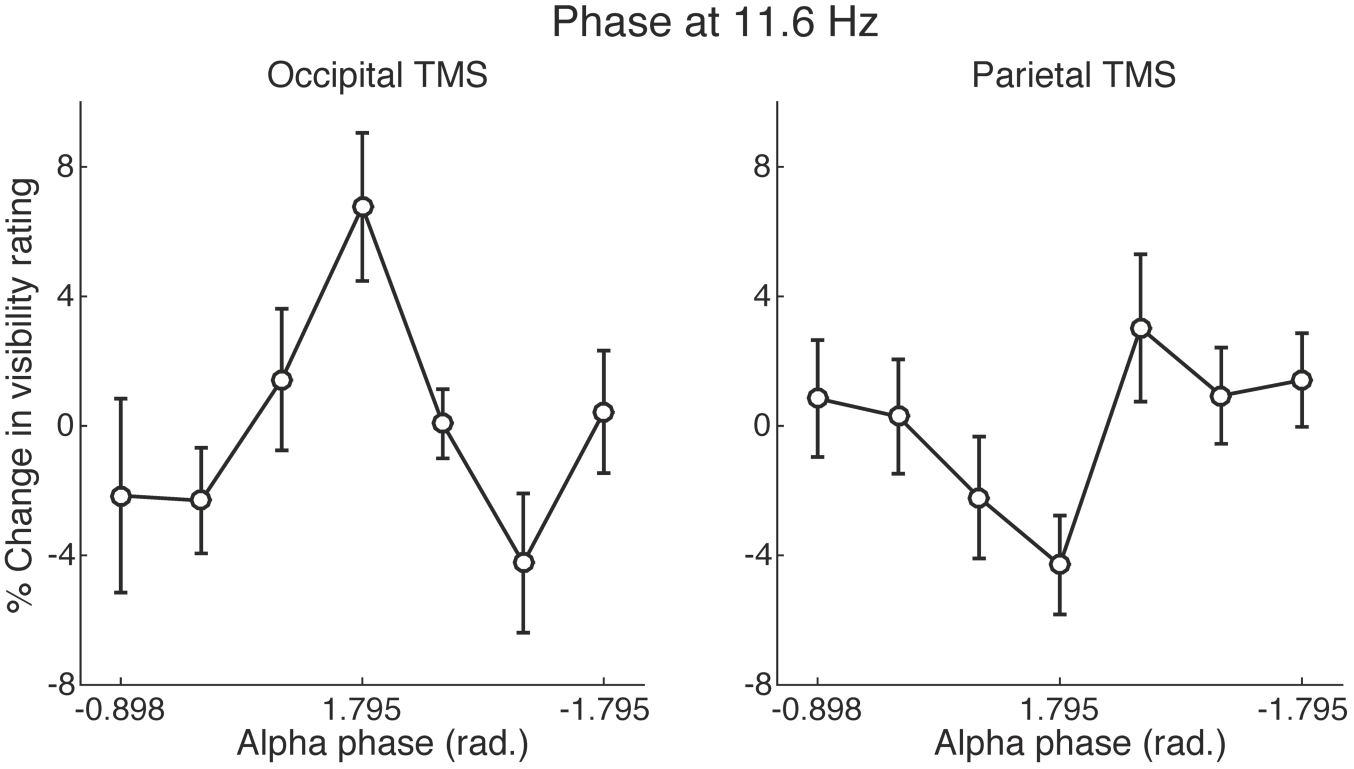
Prestimulus alpha phase predcits visibility of occipital, but not parietal TMS phosphenes. To futher inspect the relationship between prestimulus alpha phase and phopshene visiblity, we sorted trials into 7 bins according to prestimulus phase at 11.6 Hz. This revealed a significant modulation of occipital TMS phospehne visiblity of ~ 11% (peak to trough). Parietal TMS phosphenes were not significantly predicted by prestimulus 11.6 Hz phase. Error bars denote ±1 SEM.

## Discussion

We applied TMS to occipital and posterior parietal cortex at phosphene thresholds while recording EEG. This approach allowed us to investigate neural indices of cortical excitability in both regions and to track activity related to occipital and parietal TMS phosphene perception. We replicated two previous findings demonstrating a negative relationship between prestimulus alpha power and phosphene reports (4) as well as a dependence of phosphene perception on prestimulus alpha phase (10). By investigating a range of frequencies we show that prestimulus power in the delta/theta range (2-8 Hz) is also predictive of occipital TMS phosphenes. Notably, when phosphenes were induced through stimulation of PPC, we observed, for the first time, that prestimulus beta-band power (13-20 Hz) is negatively correlated with phosphene visibility. No phase-dependence was observed. TMS-evoked responses differentiated levels of awareness ~200 ms after TMS onset at both stimulation sites and was associated with an early negativity as well as a later central parietal positivity. Time-frequency responses indicated that phosphene perception was associated with increased power in the theta range, followed by a sustained decrease of power in the alpha/beta range for both stimulation sites.

### Rethinking alpha power as a general index of cortical excitability

Prevailing theory regards alpha-band oscillations as reflecting mechanisms of cortical excitability that can be routed across brain areas according to task demands so as to suppress excitability in task-irrelevant neural populations (7, 53–57). Direct evidence linking alpha power and phase to cortical excitability comes from experiments demonstrating that phosphene perception resulting from occipital cortex TMS is predicted by alpha power (4) and phase (10) just prior to TMS onset. Likewise, the magnitude of motor-evoked potentials following motor cortex TMS is also predicted by prestimulus alpha power (5, 6) and attentional modulation of alpha power has been observed in somatosensory (58), auditory (59), and visual regions (60–62). Collectively, these findings have given rise to the notion that alpha oscillations reflect a general mechanism of functional inhibition across cortex (63). Although we replicate prior findings linking phosphene perception to prestimulus alpha dynamics during occipital TMS, our finding that prestimulus beta power was predictive of PPC TMS phosphenes directly contrasts the notion of alpha as an index of neural excitability across all of cortex. Notably, frequencies below the alpha-band were also predictive of occipital TMS phosphenes.

Several existing lines of evidence also suggest that the idea of alpha as reflecting inhibition across cortex is overly simplistic. For instance, visual-evoked multiunit and gamma-band activity in macaque inferotemporal cortex were found to be *positively* correlated with prestimulus alpha power in the local-field potential, which was also found to increase when attention was paid to visual, as opposed to auditory, input (64). Similarly, in humans, predictions about an upcoming stimulus have been shown to increase prestimulus alpha power yet result in larger evoked responses – with the two processes being positively correlated (65, 66). A possible explanation for our finding that prestimulus beta power, rather than alpha, was found to predict PPC TMS phosphenes is suggested by recent work investigating oscillatory responses to TMS of different cortical regions. This line of research found that the dominant frequency in the evoked response to TMS of occipital cortex was at ~10 Hz, whereas the dominant frequency of the PPC TMS-evoked response was at ~20 Hz (35, 36). In line with this, prestimulus beta dynamics have recently been shown to predict the global mean field amplitude of the evoked response following parietal TMS (37). Thus, beta-band oscillations may reflect the dominant oscillatory mode of PPC and alpha may reflect the dominant frequency of occipital cortex. We extend this work by showing that alpha and beta also reflect cortical excitability of early sensory cortex and PPC, respectively. Another intriguing possibility is that alpha oscillations may reflect excitability levels in primary sensory cortices, whereas higher frequency oscillations reflect excitability of higher-level regions of association cortex, including PPC.

### Neural correlates of consciousness and phosphene perception

By applying TMS at phosphene thresholds, visibility reports spanned a full range of the 100-point scale (Figure 1). This allowed us to track neural activity related to phosphene perception resulting from occipital and PPC TMS. Awareness of phosphenes from both stimulation sites was associated with enhanced VAN and LP components in the time domain (Figure 2). These two components have been intensively studied using numerous awareness manipulations of visual stimuli (for reviews, see: (20, 21, 23)), but disagreement about which reflects the neural correlate of perceptual consciousness persists (26, 28, 32). The LP potential (sometimes called the P3) has been championed as the neural correlate of consciousness by proponents of the neuronal global workspace theory (67). In this context, the LP is thought to reflect widespread activation of a network of prefrontal and parietal regions that comprise the global workspace and underlie consciousness (68). As a consequence of this proposal, consciousness is thought to emerge rather late (>300 ms) relative to the onset of a stimulus. In contrast, the VAN is typically distributed over posterior sensors, though it may have a frontal component as well (28), and is observed much earlier, typically between 180 and 260 ms (26, 27). The VAN is sometimes interpreted as reflecting local recurrent processes in visual cortex hypothesized to underlie awareness (27, 69) and support for the VAN as a correlate of consciousness has come from research suggesting that attention and introspection may account for the LP modulation with awareness (24, 25, 31), and from work demonstrating superior decodability of visibility levels from early VAN, as compared to LP potential, time-windows (70). Activity in the LP time window, however, has been shown to contain stimulus specific information that covaries with subjective awareness (67).

In the present study, both components were robustly correlated with phosphene visibility in both stimulation conditions. This finding demonstrates the similarity of electrophysiological signatures of consciousness for both artificial (TMS-induced) as well natural stimuli, and provides compelling evidence that phosphene experiences from both occipital and PPC TMS are genuine perceptual experiences. Interestingly, the LP potential modulation with awareness was very similar in topography for both stimulation conditions, being maximal over central-parietal sensors. The VAN modulation, on the other hand, displayed a qualitatively different topography, being more pronounced over lateral temporal-occipital sensors following occipital TMS, and having a central occipital-parietal distribution following PPC TMS. This pattern closely resembles that observed in a recent study examining ERPs to occipital and parietal TMS phosphene perception (17). The authors found that occipital TMS phosphenes were associated with early modulation over occipital-temporal sensors, whereas PPC TMS phosphenes were associated with early ERP modulation over parietal electrodes. The implication of these findings is that the early activity associated with phosphene perception may be generated in nearby, but distinguishable cortical regions depending on the original source of stimulation. This is intriguing because it suggests that there may be several, rather than a single, neural correlates of consciousness. It further suggests that the VAN “component” is not necessarily a unitary phenomenon reflecting the activation of a single neural region responsible for visual awareness. In fact, it has recently been demonstrated that hemianopic patients with complete loss of primary visual cortex in one hemisphere are nevertheless capable of perceiving phosphenes in their blind field if contralateral PPC is stimulated (18). This strongly undermines the presumed visual-cortical origin of PPC TMS phosphenes and supports the notion that the VAN associated with PPC TMS need not be visual-cortical in origin and need not reflect recurrent processing involving visual cortex, both of which are often assumed of the VAN component (21, 28, 70).

We also report, for the first time, oscillatory correlates of occipital and PPC TMS phosphenes. We found that phosphene visibility in both stimulation conditions was positively related to power in the theta range beginning around 140 ms and negatively related to alpha/beta power extending from 250 ms until the response screen (Figure 3). Whereas the early positive theta correlation likely reflects the same phase-locked activity underlying the VAN modulation, which occurs during the same time frame, the later negative correlation with alpha/beta power reflects dynamics that are not phase-locked with TMS onset and are thus not observed in the ERP – a signature of a truly oscillatory neural process (71). Post-stimulus alpha/beta desynchronization has long been linked to perceptual processing (72–74), and is thought to reflect disinhibition of widespread cortical networks involved in perceptual inference and decision making (33, 57). Here, we show that this well-known signature of perceptual processing extends to phosphene perception and is strikingly similar regardless of the cortical origin of stimulation.

